# Coupling of spliceosome complexity to intron diversity

**DOI:** 10.1101/2021.03.19.436190

**Authors:** Jade Sales-Lee, Daniela S. Perry, Bradley A. Bowser, Jolene K. Diedrich, Beiduo Rao, Irene Beusch, John R. Yates, Scott W. Roy, Hiten D. Madhani

## Abstract

We determined that over 40 spliceosomal proteins are conserved between many fungal species and humans but were lost during the evolution of *S. cerevisiae*, an intron-poor yeast with unusually rigid splicing signals. We analyzed null mutations in a subset of these factors, most of which had not been investigated previously, in the intron-rich yeast *Cryptococcus neoformans.* We found they govern splicing efficiency of introns with divergent spacing between intron elements. Importantly, most of these factors also suppress usage of weak nearby cryptic/alternative splice sites. Among these, orthologs of GPATCH1 and the helicase DHX35 display correlated functional signatures and copurify with each other as well as components of catalytically active spliceosomes, identifying a conserved G-patch/helicase pair that promotes splicing fidelity. We propose that a significant fraction of spliceosomal proteins in humans and most eukaryotes are involved in limiting splicing errors, potentially through kinetic proofreading mechanisms, thereby enabling greater intron diversity.

## INTRODUCTION

The splicing of mRNA precursors on the spliceosome is a signature feature of eukaryotic gene expression (Sharp, 1987). Splicing plays numerous critical regulatory roles in organisms as diverse as the unicellular budding yeast *Saccharomyces cerevisiae*, apicomplexan parasites, plants, and humans. Splicing also plays key roles in ncRNA biogenesis (Ruby et al., 2007), RNA export (Zhou et al., 2000), nonsense-mediated decay (Ishigaki et al., 2001; Le Hir et al., 2001), and genome defense (Dumesic et al., 2013). The spliceosome is a complex and dynamic assembly of small nuclear ribonucleoproteins (snRNPs) and proteins that assemble onto the intron substrate and then undergo several large rearrangements to form a catalytically active complex (Wilkinson et al., 2020). Two sequential transesterification steps mediate intron removal. Pre-mRNA splicing by the spliceosome seems surprisingly complex for a process that removes a segment of RNA from a precursor. Splicing requires eight ATP-dependent steps and about 90 proteins in *S. cerevisiae*. Much of our functional understanding of spliceosome components derives from the analysis of conditional and null mutants in *S. cerevisiae* (Wilkinson et al., 2020). Human spliceosomes appear to contain about 60 additional proteins (Wahl and Luhrmann, 2015a, b). The reason for this added complexity is not understood.

The atomic structures of the rigid core portion of the spliceosome at various stages of its cycle have been elucidated using cryoEM (Wilkinson et al., 2020). Nearly all structures have been obtained using *in vitro*-assembled spliceosomes using extracts from the budding yeast *S. cerevisiae* or from HeLa cells. While these structures have revealed that the catalytic core of the spliceosome is invariant across divergent species, proteins and structures have been identified in human spliceosomes that are not found in *S. cerevisiae* spliceosomes. While it might be imagined that the higher complexity of human spliceosomes relates to late evolutionary innovations that enabled metazoan complexity, an alternative model is that the common ancestor of *S. cerevisiae* and humans harbored a complex spliceosome, whose components were lost during the evolution of *S. cerevisiae*. There is anecdotal support for this hypothesis. For example, orthologs of a number of human splicing factors that do not exist in *S. cerevisiae* have been described in the fission yeast *Schizosaccharomyces pombe* (Chen et al., 2014; Cipakova et al., 2019; McDonald et al., 1999). Prior work indicates that the *Saccharomycotina,* the subphylum to which *S. cerevisiae* belongs, has lost introns that were present in an intron-rich ancestor, such that less than ten percent of genes harbor introns in *S. cerevisiae* (Irimia et al., 2007). As in other lineages, such loss events correlate with intron signals moving towards optimal intron signals. Thus, as introns are lost, intron signals become homogenous and lose diversity. Insofar as certain splicing factors play outsized roles in recognition of introns with divergent splice signals, such homogenization might be expected to be associated with loss of spliceosomal factors and thus overall spliceosomal simplification.

We have examined evolution of human spliceosomal protein orthologs in fungi and find that most fungal lineages, including that of the genetically tractable haploid yeast, *Cryptococcus neoformans*, encode orthologs of a large fraction of human spliceosomal proteins, indicating a complex ancestral spliceosome and substantial simplification through protein loss during the evolution of *S. cerevisiae*. For many of these proteins, we have obtained viable gene deletion strains of *C. neoformans* lacking an ortholog of one of these factors. Our subsequent functional studies reveal that these factors generally promote the efficiency of splicing of intron subsets that diverge in size, branchpoint-to-3’-splice site distance and intron position from average introns. Most also influence splicing decisions with a bias towards suppressing the utilization of weak/cryptic nearby 5’ and/or 3’ splice sites, indicating a role in splicing fidelity. Cryptococcal orthologs of two human proteins, GPATCH1 and DHX35, show highly correlated splicing signatures. Affinity purification-mass spectrometry (AP-MS) identification of GPATCH1-associated proteins revealed strong copurification with DHX35 as well as components characteristic of catalytically active spliceosomes. GPATCH1 and DHX35 suppress the usage of weak/cryptic nearby 5’ and 3’ splice sites, forming a G-patch/helicase pair governing spliceosomal accuracy. Orthologs of two other human G-patch-containing proteins, RBM5 and RBM17, also suppress the usage of weak nearby alternative splice sites. Their co-purified proteins indicate that they act at earlier stages of spliceosome assembly to inspect substrates. We propose that the complexity of the spliceosome enables the use of diverse introns while promoting fidelity.

## RESULTS

### Maintenance of many dozens of human spliceosomal orthologs in fungal lineages

Like the model yeasts, *S. cerevisiae* and *S. pombe*, *C. neoformans* offers a genetically tractable haploid organism in which to investigate fundamental aspects of gene expression. To emphasize the evolutionary diversity of introns in fungi, we highlight the differences between the intron sequences and abundance between *Saccharomyces cerevisiae*, *Cryptococcus neoformans* and *Homo sapiens* in Figure 1A. The *S. cerevisiae* genome is estimated to encode only 282 introns (Grate and Ares, 2002) spread over 5410 annotated genes (0.05 introns/gene), while *Cryptococcus neoformans* (H99 strain) has 6941 annotated protein coding genes harboring over 40,000 introns (Janbon et al., 2014; Loftus et al., 2005), comparable to humans (8 introns/gene), with 27,219 annotated genes and over 200,000 introns (Piovesan et al., 2019). Note that the sequences of *C. neoformans* 5’ splice sites and branchpoints are considerably more variable than those of *S. cerevisiae*, suggesting its spliceosomes, like those of humans, may be more flexible in terms of substrate utilization (Figure 1A).

**Figure 1:**
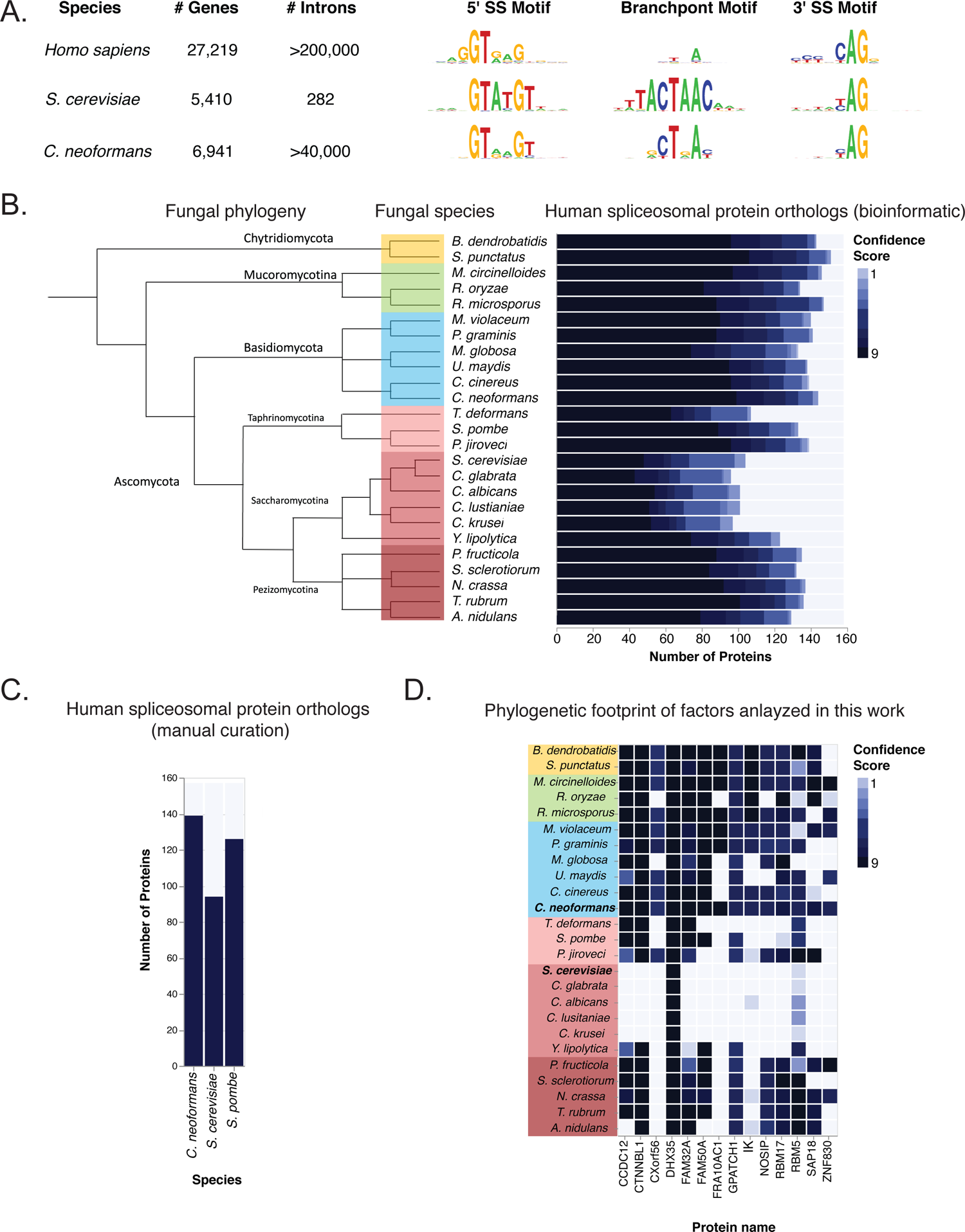
Massive loss of human spliceosomal protein orthologs in specific fungal lineages. A. Comparison of intron number and properties in humans versus the yeasts *S. cerevisiae* and *C. neoformans*. A. B. Evolutionary loss events. Shown is a phylogeny of the indicated fungal species, the groups to which they belong and the human spliceosomal protein orthologs that can be identified by a semi-automated bioinformatic process at the indicated levels of confidence (see Methods). Phylogeny is based on James et al. (2020). See also Supplementary Table S1. *B.* Numbers of human spliceosomal protein orthologs in *S. cerevisiae*, *S. pombe* and *C. neoformans*. Based on extensive additional literature curation, the numbers of human spliceosomal orthologs in the indicated species are plotted. See also Supplementary Table S1. C. Spliceosomal factor orthologs for which null mutations in *C. neoformans* were obtained. Plotted are the confidence scores for the presence of the indicated human spliceosomal protein orthologs in the indicated species. See also Supplementary Table S1.

We asked whether the loss of introns in the *Saccharomycotina*, the subphylum in which *S. cerevisiae* belongs, is accompanied with a loss of spliceosomal protein orthologs. We compiled a list of all spliceosome components reproducibly detected through mass spectrometry, interaction studies and/or purified and visualized in the spliceosome in structural biology studies (Cvitkovic and Jurica, 2013; Wahl and Luhrmann, 2015a, b). This list includes 157 human proteins (Supplementary Table S1). To identify candidates for fungal orthologs we used a combination of criteria including reciprocal BLASTP searches and the presence of predicted protein domains, followed by the application of additional criteria. Because ortholog identification is not an unambiguous exercise, we generated a confidence score (0-9) for the presence of an ortholog in given species (see Methods). Using this semi-automated process, we analyzed 24 fungal species with at least two representatives from each major clade (Figure 1B, left panel). We then plotted the number of proteins for which an ortholog to a human spliceosomal protein could be identified at a given confidence level in each species (Figure 1B, right panel). Strikingly, members of the intron-reduced *Saccharomycotina* harbored the fewest strong human spliceosomal protein orthologs. Other species exhibited considerably larger numbers of human spliceosomal orthologs, including *C. neoformans* (Figure 1B right panel). Because members of the most early branching groups analyzed harbor the highest number human spliceosomal protein orthologs, clades displaying lower numbers of orthologs have most likely undergone gene loss events, with the *Saccharomycotina* exhibiting the highest degree of loss. This correlates with the reduction in intron number found in species of this group (Irimia and Roy, 2008). For three fungal species of interest (*S. cerevisiae*, *S. pombe* and *C. neoformans*), we performed detailed manual curation of the spliceosome using the literature (including our past studies of purified Cryptococcal spliceosomes – Burke et al, 2018) and an available experimentally curated database (Cvitkovic and Jurica, 2013). Nine proteins in *S. cerevisiae* and one protein in *S. pombe* are included in the curation based on the literature despite the fact they display insufficient sequence identity with the presumptive human ortholog to be detected bioinformatically. This analysis revealed 94 spliceosomal protein orthologs in *S. cerevisiae*, 126 in *S. pombe* and 139 in *C. neoformans* (Figure 1C and Supplementary Table S1).

Thus, some 45 genes encoding predicted human spliceosomal orthologs are present in *C. neoformans* but not in *S. cerevisiae.* To investigate these spliceosomal proteins, we searched for viable knockout mutants in these factors in a gene deletion collection for *C. neoformans* constructed in our laboratory and identified strains deleted in 13 of these putative spliceosomal factors. We also identified a strain harboring a deletion of an ortholog of the human spliceosomal protein DHX35, which is found in *S. cerevisiae* (Dhr2) but appears to not be involved in splicing but instead nucleolar ribosomal RNA processing (Colley et al., 2000). We identified the Cryptococcal ortholog of DHX35 previously in purified *C. neoformans* spliceosomes (Burke et al., 2018), and thus included it in this study. The names and confidence scores in fungi of these 14 human spliceosomal proteins are displayed in Figure 1D. For readability, we will use the human nomenclature for Cryptococcal spliceosome proteins throughout the manuscript (see Supplementary Table S1 for *C. neoformans* gene locus and name). We also identified a viable gene deletion corresponding to Rrp6, a nuclear exosome subunit, involved in RNA degradation and quality control, whose loss we hypothesized might stabilize RNAs produced by aberrant splicing events compared to wild-type cells.

To examine the impact of these 14 gene deletion mutations on the abundance of pre-mRNA and mRNA along with splice site choice, we cultured these strains, extracted RNA, purified polyadenylated transcripts, and performed RNA-seq. To maximize the quality of the data, the samples were grown in duplicate and paired with duplicate wild-type samples grown on the same day to the same optical density. In addition, paired-end 100 nt reads were obtained at a minimum depth of 12M reads/sample.

### Limited impact of spliceosomal protein mutations on global transcript abundance

We first sought to determine whether deletions of putative spliceosomal proteins altered the transcript levels of other spliceosomal proteins. Hence, we subjected RNA-seq reads to mapping and applied DESeq2 to identify changes in transcript levels (Love et al., 2014). Shown in Figure 2D is the impact of gene deletions on the levels of spliceosomal protein-encoding transcripts (see Supplementary Table S2 for full results). Among the deletions analyzed only one displayed a significant change (>2 fold change, adjusted p-value<0.01) in the transcript levels of a spliceosome-encoding protein. This strain is deleted for *CNAG_02260,* which encodes the Cryptococcal ortholog of FAM50A. While FAM50A is a spliceosomal protein in humans, it has also been linked to transcription (Kim et al., 2018) suggesting a pleiotropic role. Consistent with this, we observed that many genes display transcript level changes in this mutant, while few global transcript changes were observed for the other gene deletion strains, save for the Δ*rrp6* strain, which increased the levels of ∼250 mRNAs, consistent with its predicted role in nuclear RNA turnover (Figure 2D). Thus, mutations in putative spliceosomal factors analyzed here do not generally appear to have large effects on the expression of other spliceosomal factors, suggesting that effects on splicing in the corresponding *C. neoformans* mutants likely reflect direct roles. We therefore proceeded to analyze the impact of mutations on splicing.

**Figure 2:**
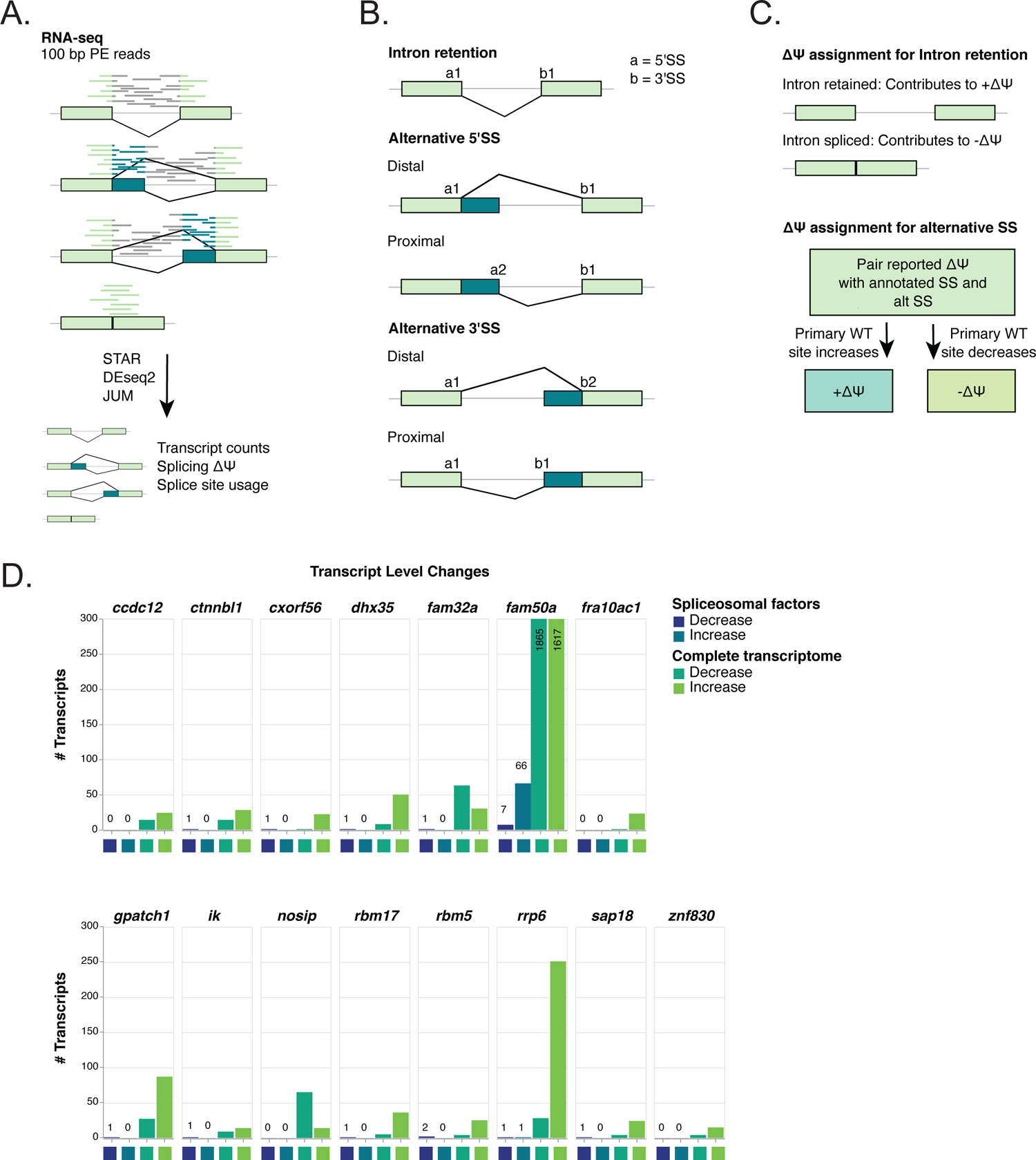
RNA-seq analysis of null mutations in 14 human spliceosomal protein orthologs. A. Schematic of the RNA-seq pipeline. Replicate mutant and paired replicate wild-type samples grown on the same day were subjected to 100nt paired-end stranded RNA-seq analysis. Initially, an annotation-free method is used to identify splicing events which was then filtered to require that at least one of the alternative events corresponded to an annotated intron. This approach produced results that could be hand-validated through direct visualization of read data. B. Definitions of splicing events quantified. Definitions of changes to Percent Spliced In (delta PSI or Δψ) values for intron retention, alternative 3’ splice site, and alternative 5’ spice site events. For the intron retention category, introns that display an increased in unspliced precursor (relative to spliced mRNA) in the mutant genotype are assigned a positive Δ C. ψ, while those that show a relative decrease in unspliced precursor are assigned a negative Δψ. For alternative 3’ and 5’ events, the primary splice site used in wild type is used to assign direction of the splicing change. An increase in primary site usage was assigned positive Δψ, and a decrease was assigned negative Δψ. D. Changes in transcript levels in mutants. 100 bp paired-end RNA-seq results for 15 knockout strains and KN99 wildtype were analyzed using DEseq2 with a 2-fold change cutoff and an adjusted p-value cutoff of 0.01. All strains were grown in duplicate with duplicate wild-type strains grown on the same day. Significantly changed gene IDs were compared to a list of human splicing protein orthologs to determine the number of splicing factors affected by the KO. Plotted is the total number of splicing factors changed in the RNA-seq data as well as total transcriptome changes. (the genotype of strains with no called decreases the expression of annotated splicing factors were confirmed by direct examination of read data to show no reads corresponding to the region deleted).

### Altered splicing choice and efficiency in mutants lacking human spliceosomal protein orthologs

Prior RNA-seq analysis of splicing in wild-type *C. neoformans* strains suggests that intron retention is the major form of alternative splicing and can be altered in response to changes in environmental conditions. However, alternative 5’ and 3’ splice sites are also observed in RNA-seq data. The extent to which isoforms have distinct functions is unknown. To examine splicing changes (Fig 2A-C), we chose to use the Junctional Utilization Method [JUM; (Wang and Rio, 2018)], a high-performing annotation-free approach for measuring splicing changes in RNA-seq data that is also optimized for measuring intron retention (a surrogate for splicing efficiency). Using a stringent read-count and p-value cutoffs (see Methods), we quantified splicing changes in each of the 14 gene deletion mutants described above. Since we did not identify any instances of mutually exclusive exons and only a small handful of cassette exons, we excluded these two categories, along with the ‘complex splicing’ category, from our downstream analysis (see Methods).

As diagrammed in Figure 2B and 2C, analysis of intron retention, alternative 5’ splice site usage, and alternative 3’ splice site usage involves multiple possibilities for a mutant phenotype. For intron retention, the amount of retained intron transcripts (i.e., precursor) can be increased or decreased relative to mRNA. For the “change in Percent Splicing In” metric (Δψ), a *positive* value corresponds to an *increase* intron retention in a mutant, while a *decrease* in intron retention produces a *negative* Δψ value (Figure 2A-C). For changes in the relative use of a splice site relative to an alternative splice site, we first determined that the site preferred in wild-type cells (>50% usage relative to the alternative site) was always an annotated splice site, while the alternative site was either unannotated or annotated as an alternative site in the current *C. neoformans* H99 strain genome annotation. Whether either was proximal or distal relative to the fixed splice site was not considered. For alternative 5’ or 3’ splice site changes, a *decrease* of usage of the preferred site in mutant produces a *negative* Δψ, while an *increase* in the usage of the wild-type site in a mutant produces a *positive* Δψ value (Figure 2B-C).

For each mutant, we quantified the effects across alternative splicing events in the *C. neoformans* genome and tabulated this data across events. The results of this analysis are shown in Figure 3A (full dataset available in Supplementary Table S3). Plotted are the number of introns impacted in each gene deletion mutant for each of the three types of splicing changes. The numbers plotted above the line indicate the number of introns whose splicing is altered in a such a way to produce a positive Δψ value as defined above, while those plotted below the line represent the number of introns impacted for a given splicing type that produce a negative Δψ value as defined above. Note that because of the stringent criteria imposed on the data and limited sensitivity of RNA-seq for rare transcript classes (e.g. unspliced pre-mRNA), these numbers are likely to be substantial underestimates. We observed the largest numbers of affected introns in the intron-retention category, and the fewest in the alternative 5’ splice site category.

**Figure 3:**
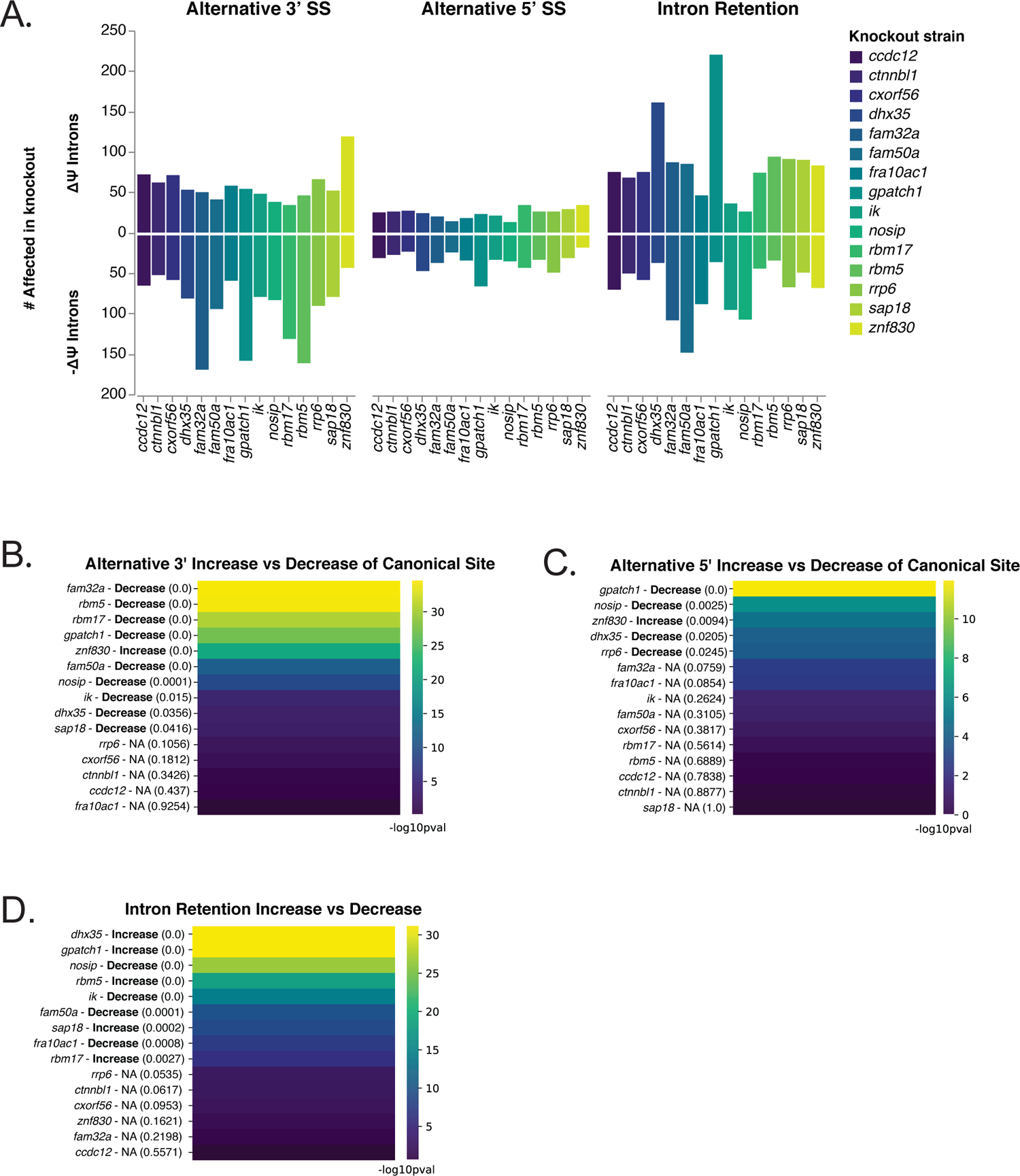
Quantification of altered pre-mRNA splicing in mutant lacking orthologs of human spliceosomal proteins. A. Number of introns altered in pre-mRNA splicing in mutants. Changes in splicing is here plotted as a count of the number of introns with significant Δψ (p<0.05) values. Counts for each KO strain are reported for each of three alternative splicing types. Intron counts displaying positive changes in Δψ as defined in Figure 2C are plotted above the line while intron counts displaying negative changes in Δψ as defined in Figure 2C are plotted below the line. Binomial test for directionality of alternative 3’ splice site usage changes. Introns affected by each KO strain were analyzed to test for a bias towards positive or negative Δ A. B. ψ. KO name is reported followed by direction and p-value. -log_10_(p-value) is displayed and colored as indicated. The labels on the left indicate the mutant, whether the bias reflects a decrease or increase in the canonical splice site in the mutant with the p-value shown in parenthesis. B. Binomial test for directionality of alternative 5’ splice site usage changes. Introns affected by each KO strain were analyzed to test for a bias towards positive or negative Δψ. KO name is reported followed by direction and p-value. -log_10_(p-value) is displayed and colored as indicated. The labels on the left indicate the mutant, whether the bias reflects a decrease or increase in the canonical splice site in the mutant with the p-value shown in parenthesis. Binomial test for directionality of intron retention changes. Introns affected by each KO strain were analyzed to test for a bias towards positive or negative Δ A. D. ψ. KO name is reported followed by direction and p-value. -log_10_(p-value) is displayed and colored as indicated. The labels on the left indicate the mutant, whether the bias reflects a decrease or increase precursor accumulation in the mutant with the p-value shown in parenthesis.

It appeared that many of the mutants were biased towards a negative Δψ for 3’ and 5’ splice site choice, indicating a decrease in the use of the canonical splice site in the mutant (and therefore an increase in the use of an alternative site). Likewise, for intron retention, several mutants appeared to be biased towards increasing intron retention, consistent with increased splicing defects (increased pre-mRNA vs. mRNA). To test the statistical significance of these apparent skews, we used the binomial distribution to model the null hypothesis. As can be seen in Figure 3B, nine deletion mutants displayed statistically significant bias towards decreased usage of the canonical site (and therefore increased use of an alternative site) for 3’ splice site usage. These correspond to strains lacking orthologs of human FAM32A, RBM5, RBM17, GPATCH1, FAM50A, NOSIP, IK, DHX35 and SAP18 (note that in humans RBM5 and RBM10 are paralogs; we refer to the Cryptococcal ortholog as RBM5 for simplicity). Curiously, a mutant lacking the ortholog of ZNF830, a human spliceosomal protein of unknown function, displayed a bias towards increased use of the canonical 3’ splice site. For alternative 5’ splice site usage, we observed a similar pattern, with cells lacking orthologs of GPATCH1, NOSIP, and DHX35 displaying a bias towards decreased use of the canonical 5’ splice site and increased use of an alternative 5’ splice site in the mutant (Figure 3C). Again, cells lacking ZNF830 displayed the opposite bias. Finally, five mutants displayed a bias towards an increase in intron retention in the mutant (DHX35, GPATCH1, RBM5, SAP18 and RBM17) suggesting a role in splicing efficiency for a subset of transcripts (Figure 3D). Unexpectedly, strains lacking orthologs of NOSIP, IK, FAM50A, and FRA10AC1, human spliceosomal protein of unknown function, display reduced intron retention in the mutant, indicating that their absence results in increased splicing efficiency for a subset of transcripts, an unexpected phenotype (Figure 3D). This initial analysis reveals categories of splicing changes produced by mutations in the factors analyzed and statistically significant biases in the directionality of these changes.

### Clustering of gene deletions based on splicing changes suggests some factors act together

To identify candidates for spliceosomal factors that might act together, we calculated correlations of splicing effects for each pair of factors. Specifically, we calculated vectors of log_10_ corrected P-values (produced by JUM’s linear model approach) for each of the ∼40,000 *C. neoformans* introns, with nonsignificant p-values corrected to 1; and then calculated correlations for these vectors for each pair of mutants. Figure 4 displays these correlation matrices for three types of splicing events using a Pearson correlation as the distance metric. Strikingly, we observed that GPATCH1 and DHX35 were nearest neighbors in all three matrices, suggesting they consistently impacted overlapping intron sets. We also noticed that RBM5 and RBM17 were also nearest neighbors in the autocorrelation matrix for the alternative 5’ splice site usage data and the intron retention data (Figure 4B-C). Human GPATCH1 and DHX35 have both been identified in C complex spliceosomes assembled *in vitro* (Ilagan et al., 2013), while RBM5 and RBM17 have been found in the early A complex that includes U2 snRNP (Hartmuth et al., 2002). Loss of the nuclear exosome factor Rrp6 produced a signature that tended to cluster adjacent to that of strains lacking the ortholog of CTNNBL1 (Figure 3A-C), a core component of active human spliceosomes recently visualized by cryoEM (Townsend et al., 2020), indicating an overlap between RNA species normally degraded by Rrp6 and those that accumulate in cells lacking CTNNBL1. Other mutants also showed some degree of clustering, suggesting functional/biochemical relationships.

**Figure 4:**
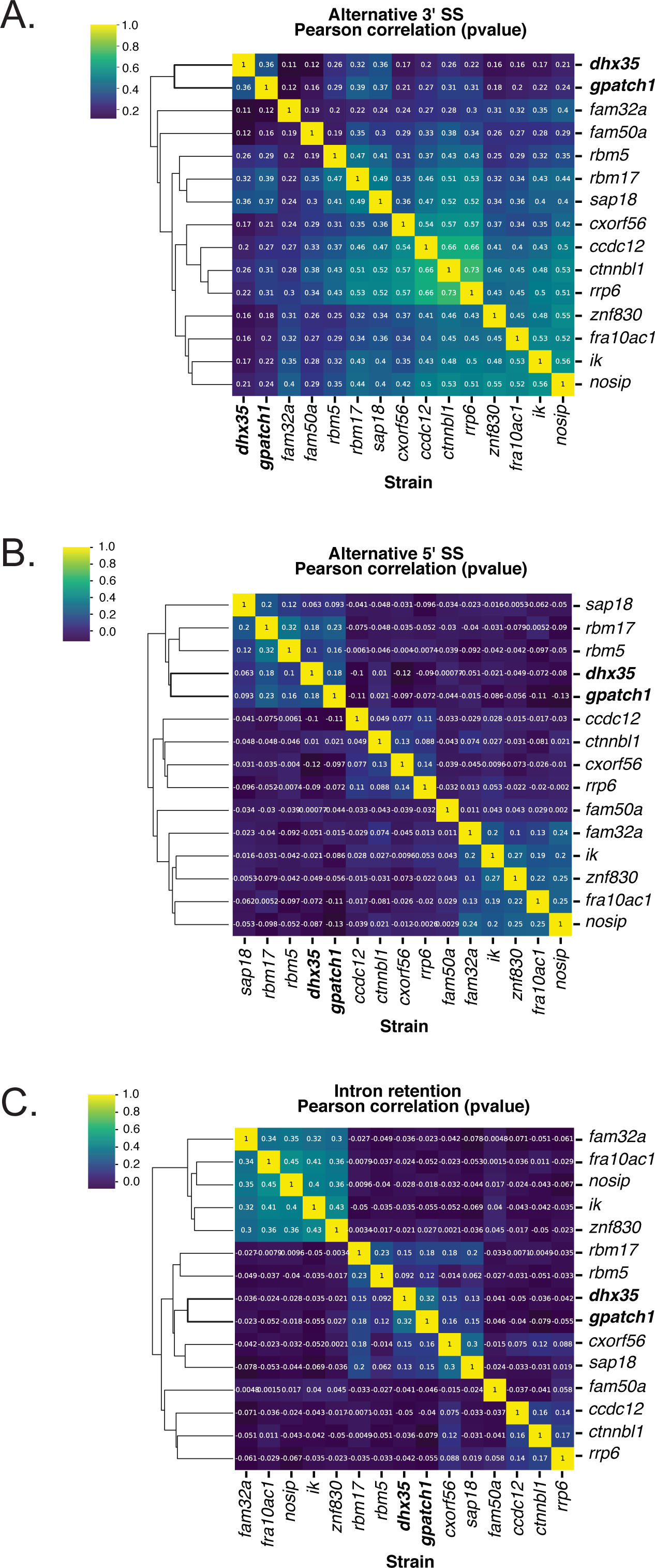
Correlation between phenotypic signatures of spliceosomal protein ortholog gene deletion mutants. P-values (corrected for multiple hypothesis testing) for changes in splicing were treated as vectors and used to generate an autocorrelation matrix for each type of splicing event. P-values greater than 0.05 were set to 1. Pearson correlation was used as the distance metric (values shown in boxes – color bar shows absolute values). Data are organized by hierarchical clustering. A. Mutant autocorrelation matrix based on significant alternative 3’ splice site changes B. Mutant autocorrelation matrix based on significant alternative 5’ splice site changes C. Mutant autocorrelation matrix based on significant intron retention changes

### GPATCH1 and DHX35 as well as RBM5 and RBM17 associate in spliceosomes

The genetic data above together with existing data on the association of the human orthologs suggests that GPATCH1 and DHX35 might act together. This would require for them to be present in the same spliceosomal complex(es). To test this hypothesis, we generated a FLAG-tagged allele of GPATCH1 and performed immunoprecipitation (IP) of an untagged strain and of the tagged strain under low and high-salt conditions (four IPs total). To quantify the proteins in the coimmunoprecipitated material we performed tandem mass tag (TMT) mass spectrometry analysis (Figure 5A). We then ranked proteins based on relative peptide counts/protein length for all identified proteins. Remarkably, the next most abundant protein in the GPATCH1 IP was DHX35 (Figure 5B). Numerous additional spliceosomal proteins were identified, including those characteristic of active C complex spliceosomes (Figure 5B), suggesting that, as in human cells, GPATCH1 and DHX35 associate with active spliceosomes in *C. neoformans*. The full mass spectrometry dataset can be found in Supplementary Table S4, which includes raw and annotated data.

**Figure 5:**
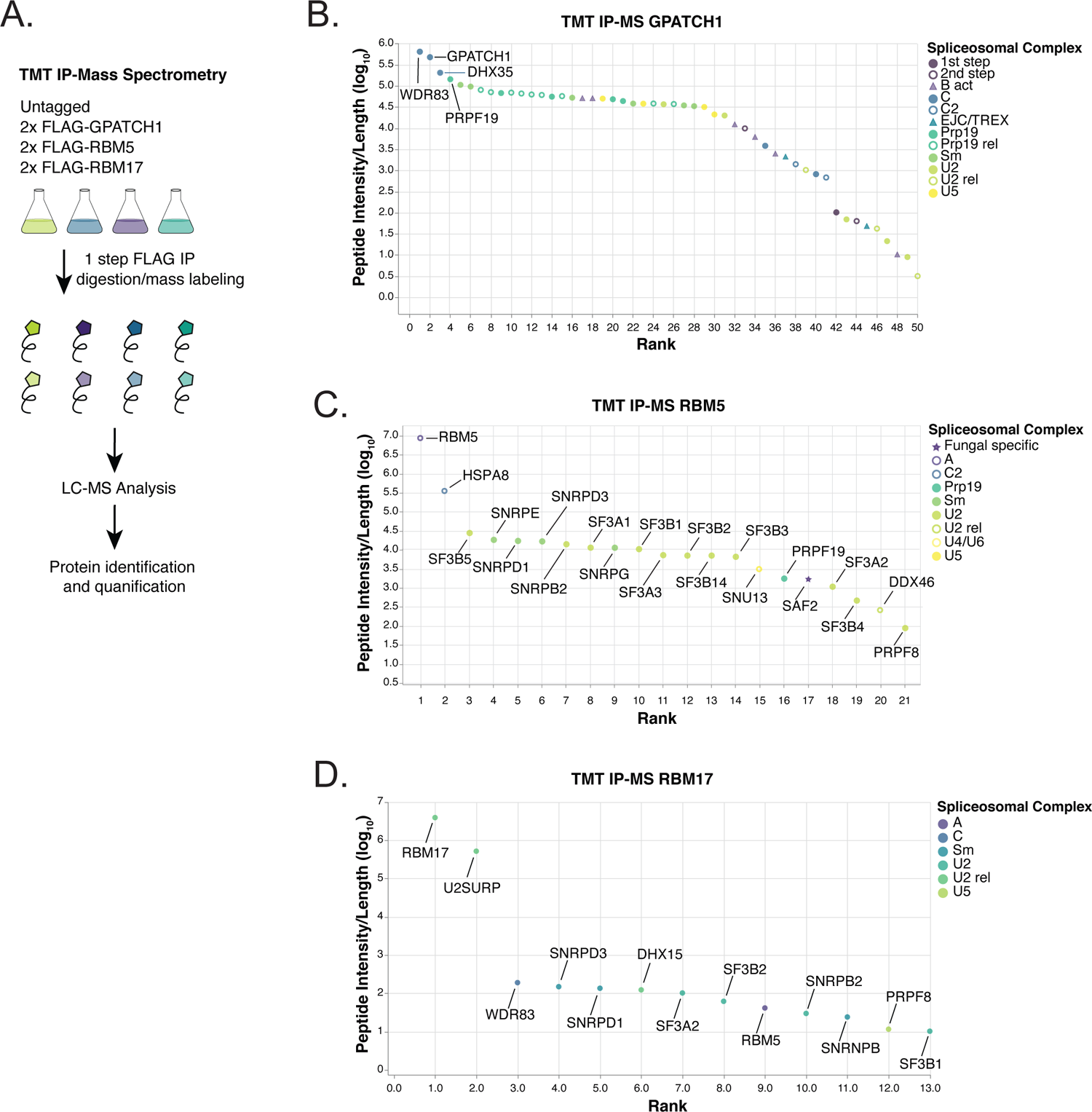
Purifications and TMT-MS analysis of endogenously tagged human spliceosomal protein orthologs. A. Schematic of sample preparation for TMT-MS. Strains harboring 2x FLAG-tagged alleles were grown to OD600nm of 2 before being harvested and lysed (see Methods). Purifications were performed on the soluble fraction in untagged and tagged strains at low and high salt (four purifications for each tagged strains). Shown are relative normalized abundances of the sum of the low- and high-salt peptide intensities of the spliceosomal protein orthologs. B. IP-MS results for 2x-FLAG GPATCH1. Plotted are TMT-MS data with length-normalized peptide intensity (log_10_) on the Y-axis and rank on the X-axis. Color and shape indicate the spliceosomal complex associated with the human orthologue. The top four hits are labeled with their orthologous human protein names. C. IP-MS results for 2x-FLAG RBM17. Plotted are TMT-MS data with length-normalized peptide intensity (log_10_) on the Y-axis and rank on the X-axis. Color and shape indicate the spliceosomal complex associated with the human orthologue. All hits are labeled with their orthologous human protein names. D. IP-MS results for 2x-FLAG RBM5. Plotted are TMT-MS data with length-normalized peptide on the Y-axis (log_10_) and rank on the X-axis. Color and shape indicate the spliceosomal complex associated with the human orthologue. All hits are labeled with their orthologous human protein names.

We also performed parallel IP experiments with RBM5 and RBM17 (eight additional purifications), as they also harbor a G-patch domain and displayed clustering in their functional signatures. These proteins displayed different associated proteins. RBM5 associated with components of the U2 snRNP including DDX46 (*S. cerevisiae* (*Sc.*) Prp5), SF3A3 (*Sc.* Prp9) and SF3A2 (*Sc.* Prp11) along with SF3B complex proteins (Figure 5C), consistent with its association with A complex spliceosomes during *in vitro* splicing reactions (Hartmuth et al., 2002). RBM17 has been found to associate with U2SURP and CHERP in IP-MS studies from human cells (De Maio et al., 2018). Strikingly, we found that purification of *C. neoformans* RBM17 identified U2SURP as the most abundant coimmunoprecipitating protein (Figure 5B), indicating evolutionary conservation of this association. We also identified peptides corresponding to RBM5 (Figure 5D), consistent with their clustering in the autocorrelation matrix based on the RNA-seq data described above. The full mass spectrometry datasets for the RBM5 and RBM17 purifications can be found in Supplementary Tables S5 and S6. Taken together, these data indicate that the clustering of factors based on their impact on splicing choice and efficiency is a useful way to begin to understand their biochemical relationships in the spliceosome.

### Identification of intron features that correlate with sensitivity to dependence on specific factors

To investigate why some introns are sensitive to loss of the spliceosomal protein orthologs described above, we tested whether 5’ splice site strength, predicted branchpoint strength (see Methods), or 3’ splice site strength was distinct for introns affected in each of the mutants studied. These studies identified only weak or marginal effects. Next, we investigated intron geometry. Specifically, we asked whether the intron length distributions of affected versus unaffected introns differed (as determined by a corrected Wilcoxon Rank Sum Test) for a given mutant and splicing type. We performed the same for the number of intronic nucleotides between the predicted branchpoint and the 3’ splice site. All mutants that impacted the splicing of introns skewed significantly towards affecting introns with longer lengths (Figure 6A; clustered heatmap of corrected p-values is shown on the left panel and top three mutants/splicing types are shown in cumulative density plots on the right). The impact was strongest for intron retention changes (Figure 6A). Differences in branchpoint-to-3’ splice site distance [both increases and decreases; denoted by (+) and (-)] were most notable of introns affected for intron retention for RBM17, CCDC12, FAM32A, and FAM50A. FAM32A has been identified a “metazoan-specific” alternative step 2 factor in human spliceosome cryoEM structures that promotes the splicing *in vitro* of an adenovirus substrate, harboring a relatively short branchpoint-to-3’ splice site distance (Fica et al., 2019). The analysis described here suggests it also limits the splicing of longer introns as well as those with nonoptimal branchpoint-3’ splice sites *in vivo*. These analyses indicate that intron size and branchpoint-3’ splice site distance are significant determinants of sensitivity to loss of the spliceosomal proteins investigated here.

**Figure 6:**
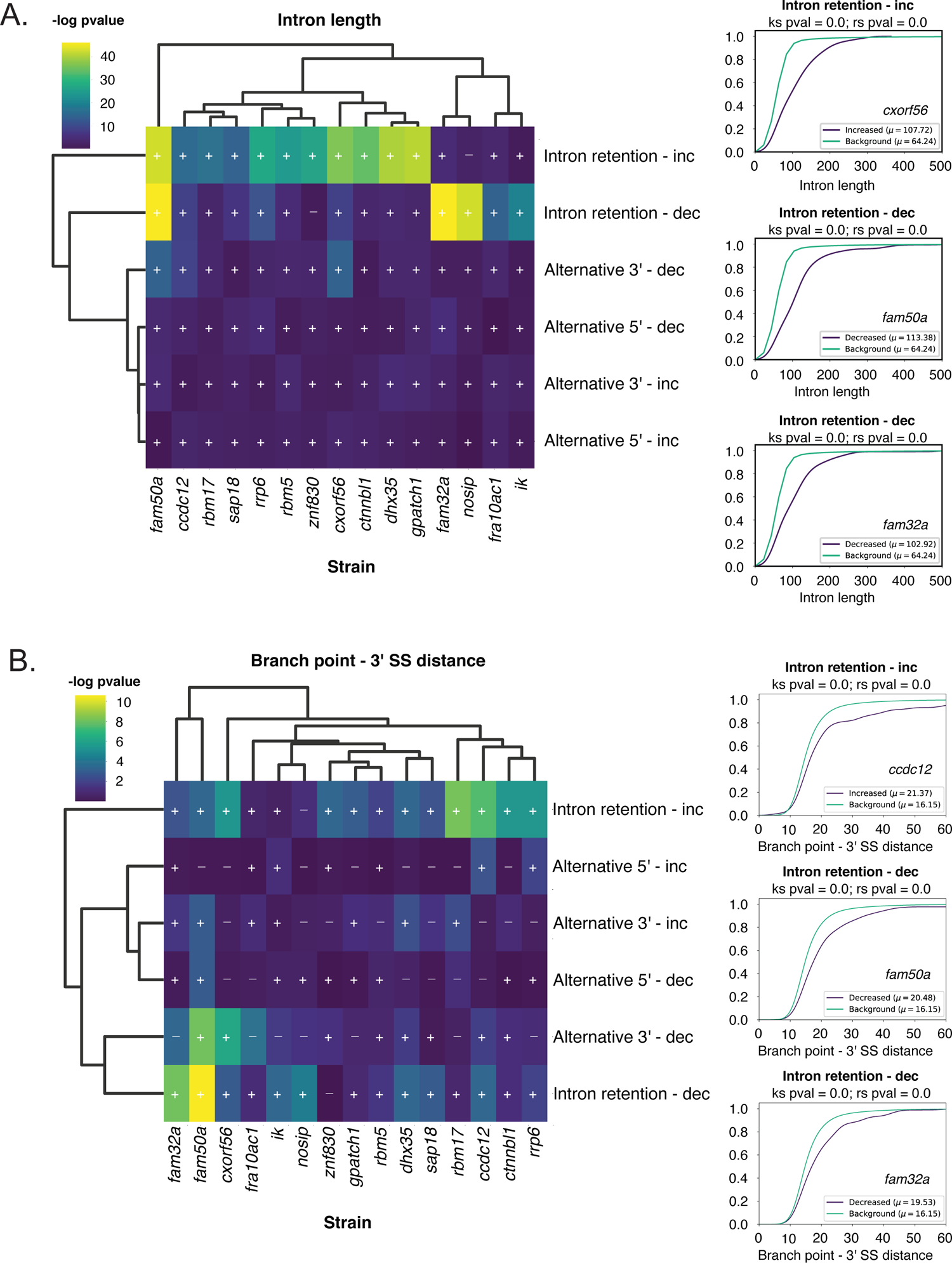
Enrichment of divergent geometry parameters of introns whose splicing is altered in human spliceosomal ortholog mutants. A. Enrichment of altered lengths in affected introns. Shown in the heatmap is the negative log of the p-value generated produced by a Wilcoxon Rank Sum Test (corrected for multiple hypothesis testing) comparing affected and unaffected introns for a given gene deletion strain and type of splicing change. Also indicated by a (+) or (-) sign is the direction of effect. Shown on the right are CDF plots and statistical test results for three gene deletion mutants/splicing change combinations that display the most significant effects. B. Enrichment of altered predicted branchpoint-to-3’ splice site distances. Analysis was performed as in A. Branchpoint-to-3’ splice site distances were predicted by using *C. neoformans* branchpoint consensus to predict branchpoints computationally.

### Mutants result in activation of weak alternative 5’ and/or 3’ splice sites

As many mutants that we examined were found to trigger reduced use of the canonical 5’ or 3’ splice site and a shift toward an alternative 5’ or 3’ splice site, we asked whether the corresponding splice site sequence differed between the canonical and alternative sites. To accomplish this, we examined the frequency of each of the four bases at the first six and last six position of each intron for the canonical versus alternative 5’ or 3’ splice site. We tested whether the nucleotide biases of the canonical versus alternative site were significantly different at a given position for a given gene deletion using a corrected Chi-squared test. Plotted in Figure 7A are the results (-log_10_P value) for the first six nucleotides of the intron for the cases of alternative 5’ slice site usage. We observed highly significant differences at many nucleotides depending on the mutant, particularly, positions 4-6 of the 5’ splice site, which normally base-pair with U6 snRNA in the spliceosome (Figure 7A). For introns displaying alternative 3’ splice site usage in the mutants, we observed significant deviation between the canonical and alternative site primarily at position −3, which is typically a pyrimidine. To examine the mutants in greater detailed, we generated sequence logo plots of the canonical and alternative sites. Shown in Figure 7C-D are those for the introns displaying decreased use of the canonical site for mutants in the three G-patch proteins analyzed above as well as DHX35. Remarkably, the alternative splice site is consistently considerably weaker than the canonical and in many cases lacking conservation at key intronic positions (e.g. positions 5 and 6 of the 5’ splice site or −3 of the 3’ splice site). We consistently observed similar patterns in mutant of the other factors. We conclude the spliceosomal proteins investigated here display a functional bias towards limiting the use of weak/alternative sites, thereby enhancing the precision of pre-mRNA splicing.

**Figure 7:**
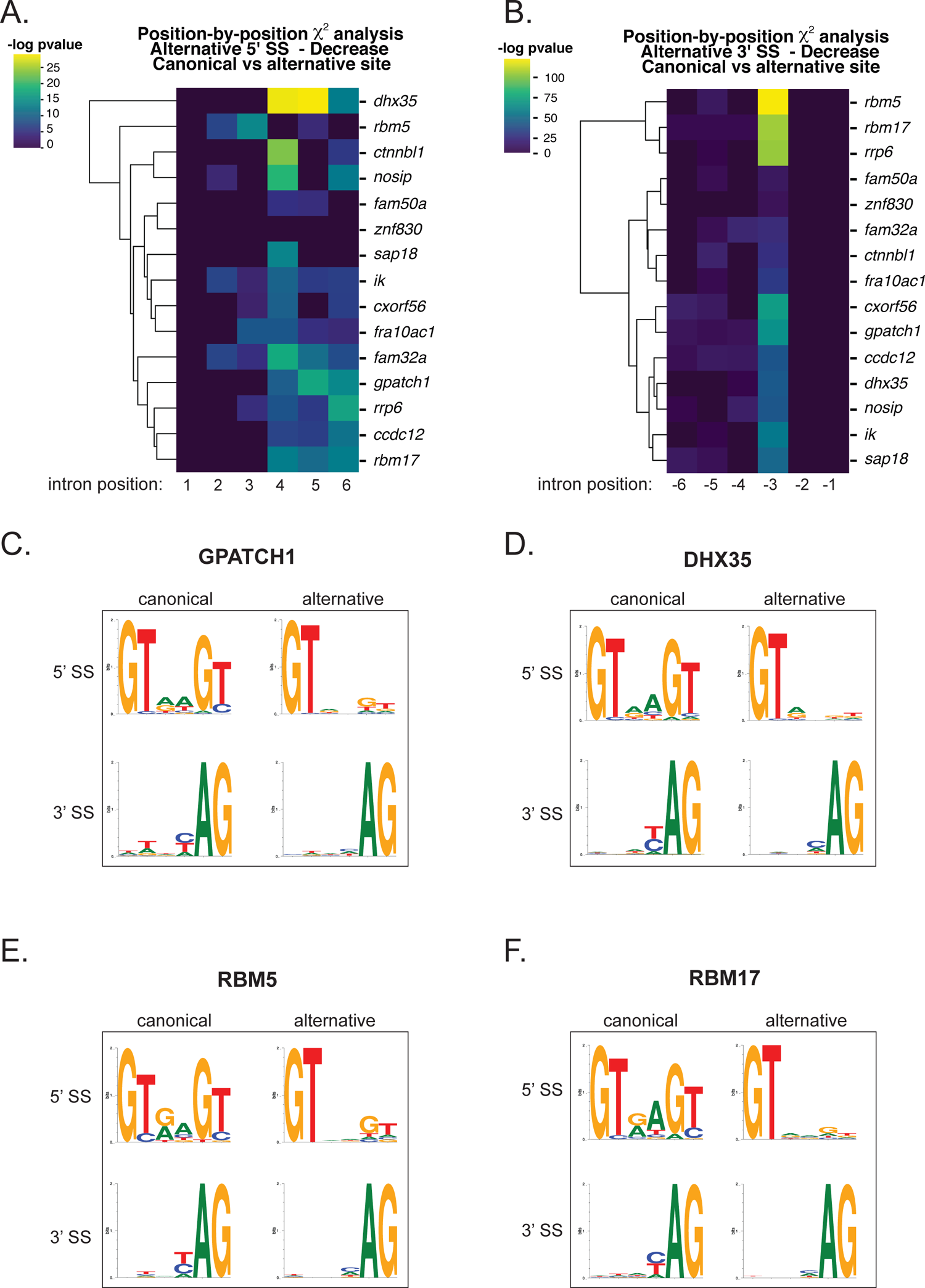
Activation of weak/cryptic alternative 5’ and 3’ splice sites in human spliceosomal ortholog mutants. 5’ splice site bases showing significant differences in composition between canonical and alternative sites. Chi-squared analysis (corrected for multiple hypothesis testing) of the first six nucleotides of introns showing significantly decreased Δ A. ψ (reduced use of the canonical site and increased use of the alternative site) for alternative 5’ splice sites in the mutant. Plotted is the negative log_10_ p value. Strains were clustered by similarity. 3’ splice site bases showing significant differences in composition between canonical and alternative sites. Chi-squared analysis (corrected for multiple hypothesis testing) of the last six nucleotides of introns showing significantly decreased Δ B. ψ (reduced use of the canonical site and increased use of the alternative site) for alternative 3’ splice sites in the mutant. Plotted is the negative log_10_ p-value. Strains were clustered by similarity.

## DISCUSSION

Our work defines a large group of spliceosomal proteins conserved between fungi and humans that enable the splicing of divergent introns while promoting fidelity. Most of these proteins have not been investigated functionally *in vivo* in any system. These factors are not essential for splicing *per se* as they were lost in large numbers during the evolution of intron-reduced species. Nonetheless, they have been conserved at least since the evolutionary divergence of fungi and humans several hundred million years ago. In the cases investigated by immunoprecipitation and mass spectrometry, factors display biochemical interactions in *C. neoformans* that are similar to those of their human orthologs, suggesting conserved functional roles in pre-mRNA splicing. This is important because orthologs of several of the proteins investigated here are involved in disease. Mutations in FAM50A cause Armfield X-linked Intellectual Disability (Lee et al., 2020). RBM17 has been strongly implicated in the pathogenesis of Spinocerebellar Ataxia type 1 (De Maio et al., 2018; Lai et al., 2011; Lim et al., 2008; Tan et al., 2016). In addition, CXorf56 has been shown to be mutated in another inherited X-linked intellectual disability syndrome (Rocha et al., 2020; Verkerk et al., 2018). Mutations in the RBM5 paralog RBM10 cause TARP syndrome, an X-linked congenital pleiotropic developmental syndrome (Johnston et al., 2010). Finally, an inherited mutation in CTNNBL1 causes common variable immunodeficiency associated with autoimmune cytopenia (Kuhny et al., 2020). The findings discussed below provide a resource for understanding the underlying molecular pathologies of these diseases.

### Massive evolutionary loss of spliceosomal proteins in the Saccharomycotina

Prior experimental work has shown that *S. cerevisiae* spliceosomes are not very tolerant of mutations of intronic sequences away from consensus (Lesser and Guthrie, 1993), with kinetic proofreading by ATPases Prp16 (human DHX38) and Prp22 (human DHX8) limiting the splicing of mutant pre-mRNAs via discard and disassembly of substrates with kinetic defects during the catalytic stages of splicing (Koodathingal and Staley, 2013). How the spliceosomes of organisms tolerate diversity in intron splicing signals and geometries is not understood. We reasoned that spliceosomal proteins whose genes were lost during evolution of organisms undergoing intron loss/homogenization might correspond to factors and processes that promote the use of divergent introns. Our analysis suggests orthologs of about a third of human spliceosomal proteins cannot be identified in *S. cerevisiae*. However, most of these are maintained in other fungal lineages. We focused our attention on *C. neoformans*, an experimentally tractable haploid yeast whose genome is intron-rich and well-annotated. Our analysis revealed 45 genes in *C. neoformans* that encode orthologs of human spliceosomal proteins that do not appear in the *S. cerevisiae* genome. Of these, we identified 13 for which deletion alleles had been generated as part of a gene deletion effort in our laboratory. We also included the helicase DHX35 in this analysis as it is found in *C. neoformans* spliceosomes but not in those of *S. cerevisiae* (Burke et al., 2018). The human orthologs of the encoded proteins studied here associate with spliceosomes at stages ranging from early complexes such as the A complex to late catalytic/postcatalytic complexes (Cvitkovic and Jurica, 2013). Three of the proteins investigated here harbor a G-patch motif, which is found in proteins that activate superfamily 2 helicases including two involved in splicing in yeast (Robert-Paganin et al., 2015; Studer et al., 2020).

### GPATCH1 and DHX35 act together on active spliceosomes

RNA-seq analysis indicated that mutations in each of the 14 of the human spliceosome protein orthologs examined altered both splicing efficiency and choice. Clustering of the data based on the impacted introns in each mutant demonstrated that mutants lacking orthologs of GPATCH1 and DHX35 displayed consistently the most correlated signatures for multiple types of splicing changes (intron retention, alternative 5’ splice site choice, and alternative 3’ splice site choice). Affinity purification of a FLAG-tagged allele of GPATCH1 identified DHX35 as a top hit. Given that G-patch proteins are known activators of helicases, it seems very likely that GPATCH1 functions to activate DHX35 in the spliceosome, although further biochemical work will be necessary to confirm this hypothesis. What the substrate of a GPATCH1/DHX35 complex might be is unclear, but, based on the nature of the changes in splice site choice (see below), a role reminiscent to those of Prp16 and Prp22 in proofreading during the catalytic stages of splicing seems possible (Koodathingal and Staley, 2013). In this regard, we note that, in human cells, GPATCH1 and DHX35 are found in catalytically active spliceosomal complexes (Ilagan et al., 2013), and the mass spectrometry data in *Cryptococcus* presented here and elsewhere (Burke et al., 2018) indicates that this pattern of association is conserved in fungi.

### Accessory factors impact the processing of genes with divergent geometries

The mRNA-to-precursor ratio is a classic measurement of splicing efficiency (Pikielny and Rosbash, 1985; Rymond et al., 1990). As such, what is referred to as an increase in intron retention is equivalent to a decrease in mRNA splicing efficiency. A subset of mutants analyzed here display a bias in an increase in intron retention (versus a decrease), indicating a tendency towards reducing the efficiency of splicing of specific substrates when mutated, reminiscent of classic pre-mRNA splicing mutants. These include mutants lacking ortholog of GPATCH1 and DHX35 as well as mutants in NOSIP, RBM5, SAP18, and RBM17. Unexpectedly, four mutants tested either show the opposite bias (a bias towards improving splicing efficiency when absent): NOSIP, IK, FAM50A, and FRA10AC1. The effects of accessory factors on splicing efficiency correlates with distinctive features of substrates, notably longer intron size and nonoptimal predicted branchpoint-to-3’ splice site distance. We note that the impact of many of the factors studied here is biased rather than purely unidirectional. For example, while knockout of the ortholog of human GPATCH1 is strongly biased towards causing reduced use of canonical 5’ and 3’ splice sites in favor of poor alternative sites, in a minority of cases, the opposite effect is observed. This may reflect a combination of direct and indirect effects [such as competition of ‘hungry’ spliceosomes for introns (Munding et al., 2013; Talkish et al., 2019)]. Alternatively, they may represent context-dependent roles that are influenced by complex differences in intron structure and sequence. Ultimately, *in vitro* reconstitution studies (either in human or *C. neoformans* extracts) and structural studies will be required for a full mechanistic understanding.

### Accessory factors promote spliceosome fidelity

A notable finding of this work is that nine factors analyzed display functional signatures that are biased towards to the suppression of the use of nearby weak/cryptic 5’ while four factors are biased towards suppression of nearby, weak 3’ splice sites. Orthologs of GPATCH1 and DHX35 are notable in that they display this function for both 5’ and 3’ sites. This phenotype further suggests that these factors may act in a manner akin to the *S. cerevisiae* fidelity factors, Prp16 and Prp22. We propose that such an additional layer of proofreading might be necessary in organisms whose spliceosomes need to accommodate more variable intron consensus sequences as such flexible spliceosomes are likely to be more error prone. Other factors, such as the G-patch proteins RBM5 and RBM17 may have similar roles in earlier spliceosomal complexes. In this regard, we note that DHX15 (*Sc.* Prp43), the spliceosomal disassembly helicase, has consistently been observed as a component of U2 snRNP (Cvitkovic and Jurica, 2013; Hartmuth et al., 2002). Our studies suggest that much remains to be learned about the spliceosome, and points towards the key roles of functional studies in tractable non-reduced organisms in complementing studies from *S. cerevisiae*.

## METHODS

### Spliceosomal protein searches

Spliceosomal protein searches were performed on proteome assemblies available from NCBI and UniProt (See Supplementary Table S7). A curated list of relevant human spliceosomal proteins was used as queries in local BLASTp (version 2.9.0+) searches against independent Fungal proteome databases (initial e-value threshold of 10^-6^) (Altschul et al., 1997). The results from the BLAST searches were further screened by analyzing domain content (HMMsearch, HMMer 3.1b2 – default parameters), size comparisons against human protein sequence length (within 25% variation), and reciprocal best-hit BLAST searches (RBH) to the query proteome (Bork et al., 1998; Johnson et al., 2010; Tatusov et al., 1997). To avoid bias in protein domain content, domains used for HMM searches were defined as described (Hudson et al., 2019). Briefly, a conserved set of domains for each spliceosomal protein was assembled by using only those domains present in all three of the human, yeast, and *Arabidopsis* orthologs. Fungal ortholog candidates in this study were scored and awarded a confidence value of 0-9 based on passing the above criteria. Scores were calculated by starting at 9 and penalizing candidates for falling outside of the expected size range (−1 point), missing HMM domain calls (−2 points), and failing to strictly pass RBH (−5 points). A score of 0.5 was given to candidates that failed all criteria but still had BLAST hits after the initial human query to separate from those that had no BLAST hits.

### A. C. neoformans cultivation

Two-liter liquid cultures of all strains were grown in YPAD medium (Difco) by inoculation at low density (0.002-0.004 OD600 nm) followed by overnight growth with shaking 30°C. For RNA-seq experiments, cells were harvested at OD_600_ of ∼ 1. For TMT-MS experiments, an additional 1% glucose was added when the cultures reached OD_600_ of 1. Cells were harvested at OD_600_ of 2.

### Immunoprecipitation and TMT-MS

Strains harboring a C-terminal (GPATCH1 and RBM17) or an N-terminal (RBM5) CBP-2XFLAG tag were generated by homologous replacement. Immunoprecipitations were performed exactly as described (Burke et al, 2018) with the following modifications: lysis and wash buffers were adjusted to either 150 mM NaCl (low salt) or 300 mM NaCl (high salt). Two untagged samples and two tagged samples (one at each salt concentration) was produced. The four samples were then subject to TMT-MS exactly as described (Burke et al., 2018).

### RNA preparation

Polyadenylated RNA was prepared exactly as described (Burke et al., 2018).

### RNA-seq

RNA-seq libraries were prepared using the NEBNext Ultra Directional RNA Library Prep Kit for Illumina. Samples were sequenced using an Illumina HiSeq 4000 instrument. Paired-end 100 nt reads were obtained. Data are available at the NCBI GEO database: GSE168814

### RNA-seq data analysis

All reads were analyzed using FastQC and reads with more than 80% of quality scores below 25 were thrown out. Reads were aligned using STAR (Dobin et al., 2013). A minimum of 12M read/strain/replicate were obtained (Supplementary Table S8). Differentially spliced introns were called using JUM (version 2.0.2). Differential events with a p-value of greater than 0.05 were set to 1. An additional 5 read minimum was imposed. To further minimize false positive, differential splicing events called by JUM that do not have an isoform harboring a start and end corresponding with an annotated intron were also removed. Spot-checking of differential events was accomplished by manual browsing of the data. Alternative 3’ and 5’ splicing events containing more than two alternative endpoints were also removed. In few cases, JUM called introns as significantly alternatively spliced with a Δψ of 0; those introns were also removed.

Next, each intron is classified as increased or decreased and proximal or distal based on the observed canonical endpoint and associated Δψ. General data analysis, plotting and statistical testing were performed using Python and the SciPy stack as follows:

<u>Binomial tests</u>: Introns were grouped by splicing event and strain. Within each strain a binomial test (scipy.stats.binom_test) was conducted to see if there was significantly more or fewer introns with increased splicing.

<u>Comparisons of distributions</u>: Introns are grouped by strain and condition and each subset is compared to unaffected introns. The resulting two distributions are compared for each attribute. A Wilcoxon rank-sum test (scipy.stats.ranksums) is conducted to determine if the means of the two distributions are significantly different. (A Kolmogorov–Smirnov test is conducted to compare the distributions themselves (scipy.stats.kstest).) All results are multiple-test corrected using the FDR correction (statsmodels.stats.multitest.fdrcorrection).

<u>Chi-squared analysis (canonical vs alternative sites):</u> Introns were separated by strain and condition and the first and last six nucleotides of the canonical sequence of affected introns was compared to the non-canonical sequence of affected introns. Each position in the endpoints was treated as an independent Fisher exact test (FisherExact.fisherexact) or chi-square test (scipy.stats.chisquare) with (4-1)*(2-1)= 3 degrees of freedom performed on a contingency table with nucleotides in the rows and affected vs. unaffected introns as the columns. In some cases where less than 5 counts are observed in a category, a chi-squared test becomes inappropriate and the Fisher exact test is used.

<u>P-value correlations:</u> Treating all strains as a vector of p-values of affected introns, a Pearson correlation matrix is computed. (pandas.DataFrame.corr).

<u>Seqlogos:</u> Affected introns are grouped by splice type, condition (increased or decreased), and strain. Seqlogos are generated from the first and last six nucleotides (seqlogo.seqlogo).

#### SUPPLEMENTARY TABLE LEGENDS

**Supplementary Table S1:** <u>Core human spliceosomal proteins and fungal orthologs.</u> Data are presented as an Excel file.

**Supplementary Table S2:** <u>DESeq2 output of RNA-seq data.</u> Data are presented as an Excel file.

**Supplementary Table S3:** Output of Junction Utilization Package (JUM) Analysis of 14 Mutant Strains. Data are presented as an Excel file.

**Supplementary Table S4:** <u>Full TMT-MS Data for GPATCH1 immunopurifications</u>. Data are presented as an Excel file. Raw, annotated, and spliceosomal protein data displayed in separate sheets.

**Supplementary Table S5:** Full TMT-MS Data for RBM17 immunopurifications. Data are presented as an Excel file. Raw, annotated, and spliceosomal protein data displayed in separate sheets.

**Supplementary Table S6:** Full TMT-MS Data for RBM5 immunopurifications. Data are presented as an Excel file. Raw, annotated, and spliceosomal protein data displayed in separate sheets. For readability, ribosomal proteins were filtered to generate the annotated sheet.

**Supplementary Table S7:** <u>Proteomes used for Evolutionary Analysis.</u> Data are presented as an Excel file

## ACKNOWLEDGEMENTS

This work was supported by R01 GM71801 and R01 AI00272 to H.D.M. B.A.B. and S.W.R. are supported by NSF Award 1751372 to S.W.R. I.B. is supported by a Swiss National Foundation Fellowship (191929). We thank Qingqing Wang for assistance with installation and usage of JUM scripts.

